# Phylonumtomics uncovers diverse evolutionary trajectories of mitogenomic fossils buried in mammalian and avian genomes

**DOI:** 10.1101/2023.08.07.552327

**Authors:** Yu-Chi Chen, David L. J. Vendrami, Maximilian L. Huber, Luisa E. Y. Handel, Christopher R. Cooney, Joseph I. Hoffman, Toni I. Gossmann

## Abstract

Sporadically genetic material that originates from an organelle genome integrates into the nuclear genome. However it is unclear what processes maintain such an integration over longer evolutionary time. Recently it was shown that nuclear DNA of mitochondrial origin (NUMTs) may harbour genes with intact mitochondrial reading frames despite the fact that they are highly divergent to the host’s mitochondrial genome. Two major hypotheses have been put forward to explain the existence of such mitocoding nuclear genes: (A) recent introgression from another species and (B) long-term selection. To address whether these intriguing possibilities do play a role we scanned the genomes of more than 1,000 avian and mammalian species for NUMTs. We indeed identified that the subclass of divergent NUMTs harbouring mitogenes with intact reading frames are widespread across mammals and birds. We can show that for these NUMTs signatures of cross-species introgression are widespread in birds, but not mammals with the exception of ungulates. We can also show that a substantial fraction of deeply divergent NUMTs are maintained by selection. For a small number of NUMT genes we identify an evolutionary signature that is consistent with adaptive evolution including one human NUMT that is shared among seven ape species. This highlights the intriguing possibility that NUMT insertions occasionally may be functional.

## Introduction

A key question in evolutionary biology is how genetic novelties arise and contribute to adaptation (Sangster and Luksenburg 2021; Bomblies and Peichel 2022). Mutations are the ultimate source of genetic novelties that are subsequently subject to evolutionary forces such as selection and drift (Huang et al. 2016). There is compelling evidence that most mutations do not show a signature of positive selection on the molecular level, but instead are subject to random genetic drift and purifying selection (Bank et al. 2014). As novel mutations may be insufficient in many cases to generate adaptive variation, alternative mechanisms may explain the rise and manifestation of novel beneficial genetic elements.

Mutations that have already been exposed to evolutionary forces may provide a means of generating adaptive variation. For example, the acquisition of novel, pre-adapted genetic elements can occur via the lateral transfer of DNA. There is considerable evidence for such exchanges of genetic material among bacteria and even in eukaryotes (Danchin 2016; Dunning et al. 2019). For example, the substantial exchange of genetic material between eukaryotic species occurs through hybridisation and introgression (Gabaldón 2020; Setter et al. 2020). While it is known that an organism’s genomic architecture is fundamental to whether or not hybridization/introgression occurs successfully, we have limited knowledge of the extent to which lateral gene transfer, hybridisation and introgression contribute to genomic diversification driven by the acquisition of adaptive elements across species.

An intriguing possibility for the acquisition of novel, genetically adapted elements is the transfer of organelle DNA. In many eukaryotic species, organelles, such as mitochondria or plastids possess their own genomes. Mitochondrial DNA integration into the nuclear genome (**NUMT**) is a continuous and ongoing mutagenic process, which occurs across different species and tissues (Ricchetti, Tekaia, and Dujon 2004). After NUMTs were first observed in the domestic cat (*Felis catus*) (Lopez et al. 1994) they were then found in many different eukaryotic species from yeast (Ricchetti, Fairhead, and Dujon 1999) to humans (Lopez et al. 1997) and many other mammals (Tsuji et al. 2012; Calabrese et al. 2017; Gossmann et al. 2019; Biró et al. 2022; Uvizl et al. 2023) and birds (Nacer and Amaral 2017; Liang et al. 2018; Lucas, Vincent, and Eric 2022; Baltazar-Soares et al. 2023). NUMT insertion number, size and sequence composition vary not only across species (Richly and Leister 2022; Hazkani-Covo 2022) but also within populations (Soto-Calderón et al. 2014; Lucas, Vincent, and Eric 2022; Wei Wei et al. 2022). In humans, around 14% of individuals possess NUMTs that are present in less than 0.1% of the broader population and some of these NUMTs have been shown to be passed on from parents to their offspring (Wei Wei et al. 2022). This suggests that NUMT integration is a frequent mutagenic event that produces inheritable changes, potentially relevant to future evolution. The germline NUMT mutation rate in human is estimated at 2.44 × 10^−4^ mutations per generation (Wei Wei et al. 2022) illustrating that novel NUMT genetic variation is frequently and constantly being generated.

While NUMTs are widespread in the animal kingdom, there is little evidence of functional integrations of mitochondrial DNA (Noutsos et al. 2007; Pozzi and Dowling 2019; W. Wei and Chinnery 2020). Instead, it is generally assumed they are exposed to genetic drift and will quickly accumulate novel disruptive mutations. In line with this, it is frequently observed that mitocoding genes integrated into the nuclear genome accumulate missense mutations that ultimately lead to a broken reading frame. However, if the integration is very recent, the local mutation rate is very low, or selection actively maintains the coding properties of the integration, the former mitochondrial coding genes may retain intact reading frames over longer periods of evolutionary time. Such NUMTs containing genes with intact mitochondrial reading frames can be considered as **coding NUMTs (cNUMTs)**.

Recently in the genome of the Antarctic fur seal (*Arctocephalus gazella*) several cNUMTs were identified (Vendrami et al. 2022). The coding genes of three of the cNUMTs contained numerous silent mutations (i.e. showed substantial divergence at neutral sites), but very few amino-acid changing mutations relative to fur seal mitochondrial genes. The authors concluded that the signature of coding NUMTs that are divergent (**dcNUMTs**) at synonymous sites was consistent with purifying selection and hinted at functionality and introgression. As the dcNUMTs observed in the Antartic fur seal genome cannot be explained by a recent nuclear integration from it own mitochondrial genome, alternative scenarios of non-canonical integrations might explain the existence of dcNUMTs (Figure 1). Currently, it is unclear how common and frequent dcNUMTs are across larger taxonomic groups and whether they harbour signatures of functionality and introgression. Hence their potential role as adaptive elements of the genome has yet to be investigated.

**Figure 1:**
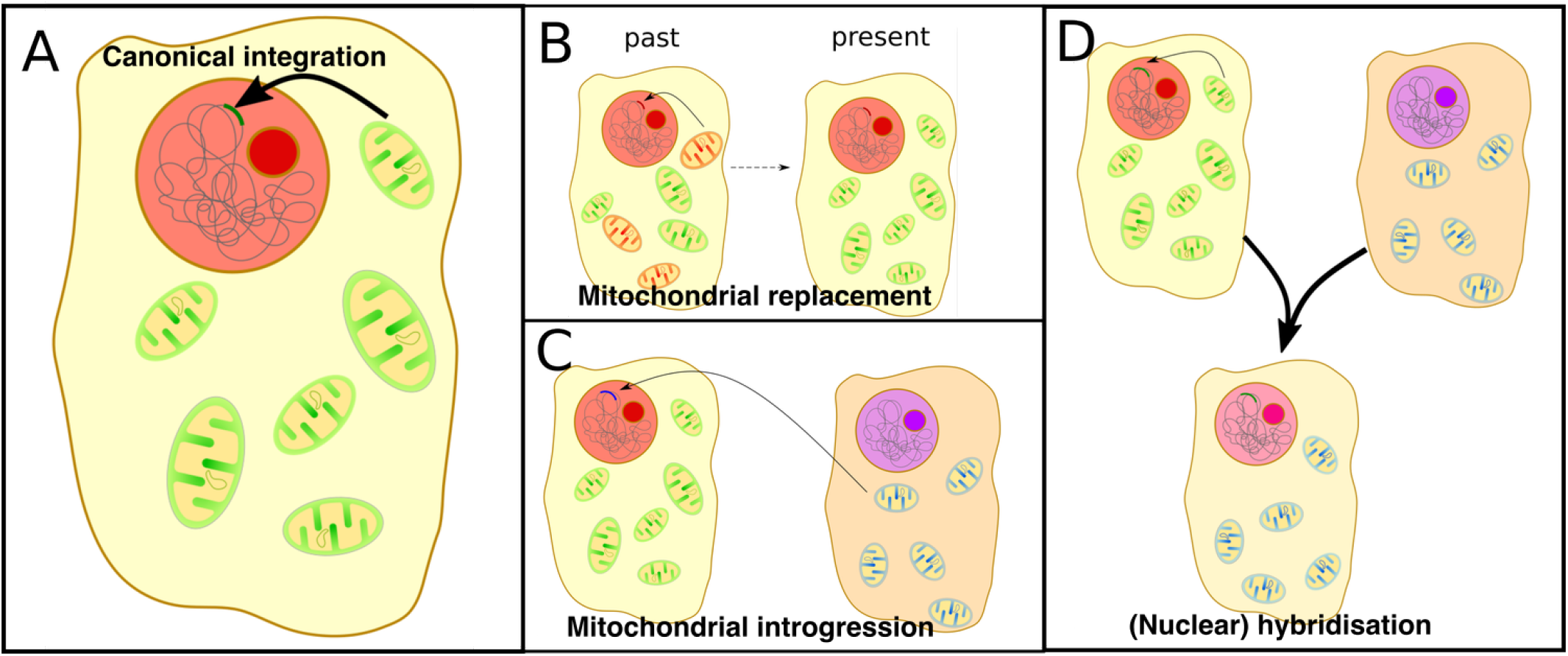
Potential sources of mitochondrial DNA of NUMTs. **(A)** Canonical integration through host species’ mitochondrial genome, **(B)** Integration of the host species’ mitochondrial genome stemming from heteroplasmic DNA, **(C)** Integration of foreign species mitochondrial DNA, e.g. through retroviruses, **(D)** Integration of nuclear DNA from another species, e.g through hybrid formation, hat happens to carry a NUMT

To unravel the role of organelle DNA in creating novel and potentially adaptive variation in nuclear genomes, we applied a comparative genomics approach, which we term ‘phylonumtomics’. We use this approach to identify NUMTs in more than 1,000 publicly available avian and mammalian genomes and to investigate the evolutionary origins of dcNUMTs. We show that dcNUMTs are widespread across mammalian and avian genomes, with distinct phylogenetic differences between mammals and birds. We conduct comprehensive analyses to uncover traces of selection by incorporating consequences of low local mutation rates and introgression. We show that in birds, the vast majority of dcNUMTs are reflects of ancient mitochondrial genomes that are preserved as ‘genomic fossils’. However, we also highlight numerous instances representing likely signatures of recent introgression, as well as a limited, but highly significant, role of positive selection. Taken together, our results suggest that mitochondria may have a much greater contribution to the evolution of the nuclear genome than previously envisaged.

## Results

### Class-wide NUMT identification

To investigate the evolutionary consequences of NUMT integration in a comparative context, we identified mitochondrial insertions into the nuclear genomes of more than one thousand mammalian and avian species. Specifically, we analysed 680 mammalian genomes and 458 avian genomes for which both a genomic and mitogenomic assembly was available from public databases. We used a previously published bioinformatic pipeline that uses the host’s mitochondrial genome DNA as a BLAST query to identify NUMT insertions in the nuclear genome (Vendrami et al. 2022). Altogether, we identify 85,100 NUMTs in 435 mammalian species (66.62%) and 28,947 NUMTS from 458 avian species (100%). This observation is in line with previous findings on 70 vertebrate species that NUMT integration is a frequent and ongoing process across mammals and birds (Hazkani-Covo 2022). We observed fewer NUMTs in birds relative to mammals (average ≈ 125 versus ≈ 63 NUMTs per species, respectively), and also find that for mammals the number of NUMTs is positively correlated with genome size, which is not the case for birds (P < 2.2 × 10^−16^ and P = 0.79, for mammals and birds, respectively, phylogenetic generalized least-squares regression).

### Phylogenetic patterns of NUMTs across mammals and birds

As we observe contrasting pattern in terms of NUMT numbers between the two taxonomic groups, we next considered the NUMT occurrences within a phylogenetic context. For this we used established species phylogenies for mammals and birds (Kumar et al. (2017), Timetree.org) and reconstructed the number of NUMTs as a phylogenetic trait using a Baysian approach with BayesTraits V4.0.1 (Pagel and Meade 2022). We observe several clear phylogenetic pattern in the data. First, the number of NUMTs varies substantially across clades (Figure 2). For example, among mammals, most marsupials show a complete lack or a very low number of NUMTs with the exception of the Tasmanian devil (*Sarcophilus harrisii*) (Hazkani-Covo 2022). In contrast, a few clades, such as old world monkeys and Cetaceans carry comparatively large numbers of NUMTs. For avian species, three clades in Passeriformes show an conspicuous excess of NUMTs. The first of these clades includes warblers, cowbirds and blackbirds. The second clade includes sparrows (e.g. *Zonotrichia albicollis*), junco, seedeaters, Tanager and finches (e.g. *Geospiza fortis*) and the third clade includes cardinals (*Cardinalis cardinalis*; Sin, Lu, and Edwards (2020)), buntings, grosbeak, longspur and grackle. Second, the number of NUMTs varies substantially between closely related species. For example for birds, *Calidris pygmaea*, a highly endangered species (Chowdhury, Foysal, and Green 2022), shows a massive number of NUMT integrations suggesting a highly degenerated genome. Mammalian species with exceptionally high numbers of NUMTs include the Tasmanian devil (*Sarcophilus harrisii*, 1,546 NUMTs), gray short-tailed opossum (*Monodelphis domestspizella passerinaica*, 811 NUMTs) and yellow-footed antechinus (*Antechinus flavipes*, 792 NUMTs) in mammals. In birds, beside the already mentioned spoon-billed sandpiper (*Calidris pygmaea*, 13,170 NUMTs) are also an island finch (*Nesospiza acunhae*, 403 NUMTs) and the northern cardinal (*Cardinalis cardinalis*, 373 NUMTs) species with a substantial NUMT enrichment in their genomes.

**Figure 2:**
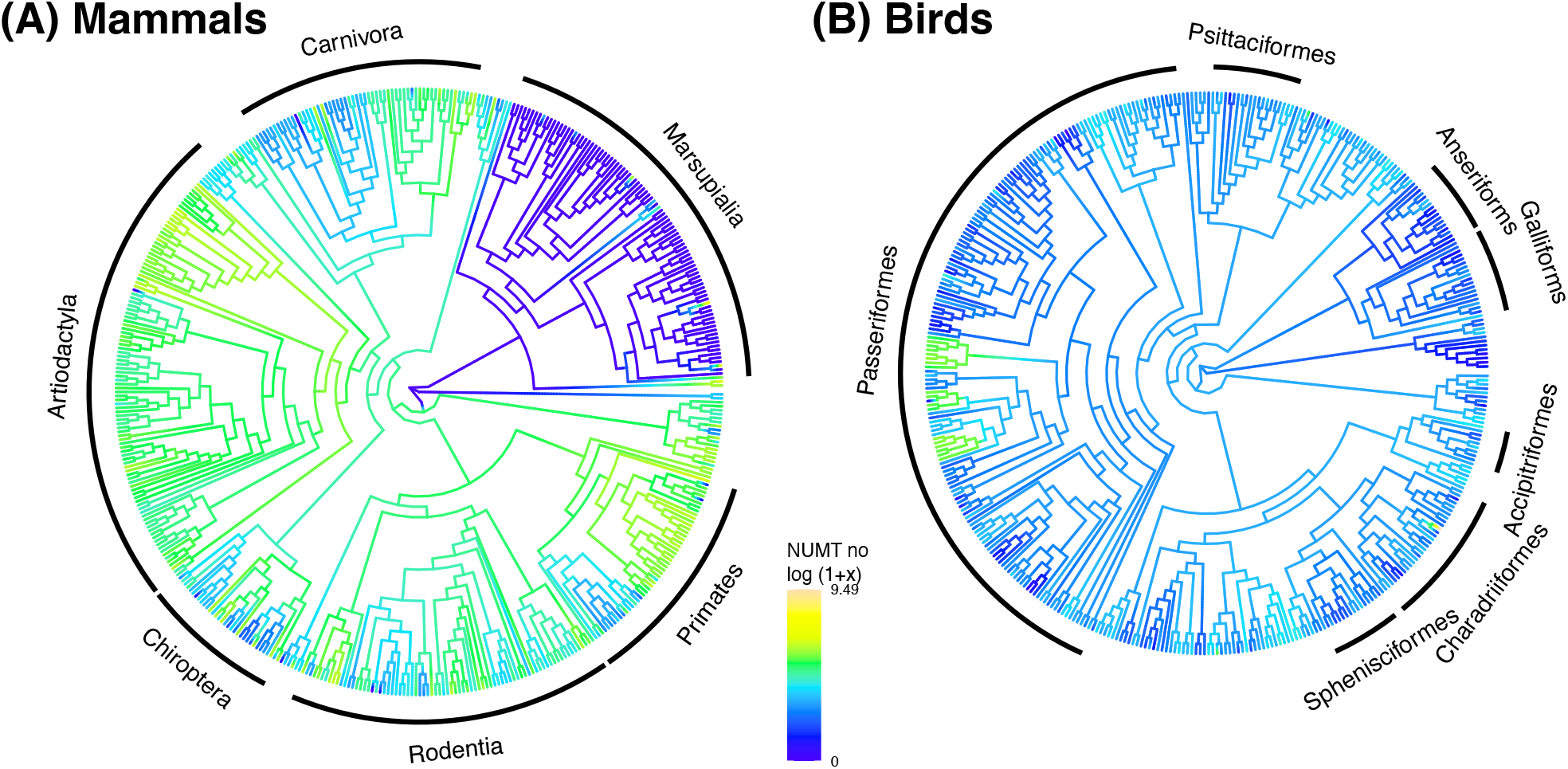
Phylogenetic distribution of NUMT occurences in mammals and birds. Shown are the species cladograms from timetree and colours represent reconstructed number of NUMTs for **(A)** mammals and **(B)** birds, respectively. Note that the colorised representation refers to the log-transformed (i.e. *log(1+x)*) NUMTs number.

### Distinct distributions of coding NUMTs, a subclass of NUMTs

While our data show that mitochondrial genome integration into the nuclear genome is a frequent and ongoing process in many species, it remains unclear what the origins and evolutionary trajectories of such NUMTs are, in particular those that are retained over long evolutionary time harbouring genes with intact mitocoding reading frames. To address this, we focus on a subgroup of NUMTs retaining at least one coding gene with an intact reading frame (coding NUMTs, cNUMTs). Among the 114,047 NUMTs identified in this study, 6,429 NUMTs from 698 species (5,229 NUMTs for 379 mammals and 1,200 NUMTs for 319 for birds) possess at least one mitochondrial gene with an intact reading frame. In mammals the number of cNUMTs is correlated with the number of NUMTs that do not have at least one gene with intact reading frame, which is not the case for birds (P < 2.2 × 10^−16^ and P = 0.9 for mammals and birds, respectively, phylogenetic generalized least-squares regression). This suggests that cNUMTs may be considered as a by-product of NUMT integration in mammals, but not in birds. Using pairwise coding divergence estimates (Wang et al. 2010), we obtain a proxy for the evolutionary age of the cNUMTs using the synonymous divergence between the cNUMT genes and the respective genes on the host mitochondrial DNA. The distribution of these *dS* values may uncover common integration times across species (Figure 3). We find that while the number of cNUMT becomes lower for increasing *dS* values in mammals, for birds there is a clear peak at *dS* ≈ 1, which might reflect a historic episode of high transposable element activity.

**Figure 3:**
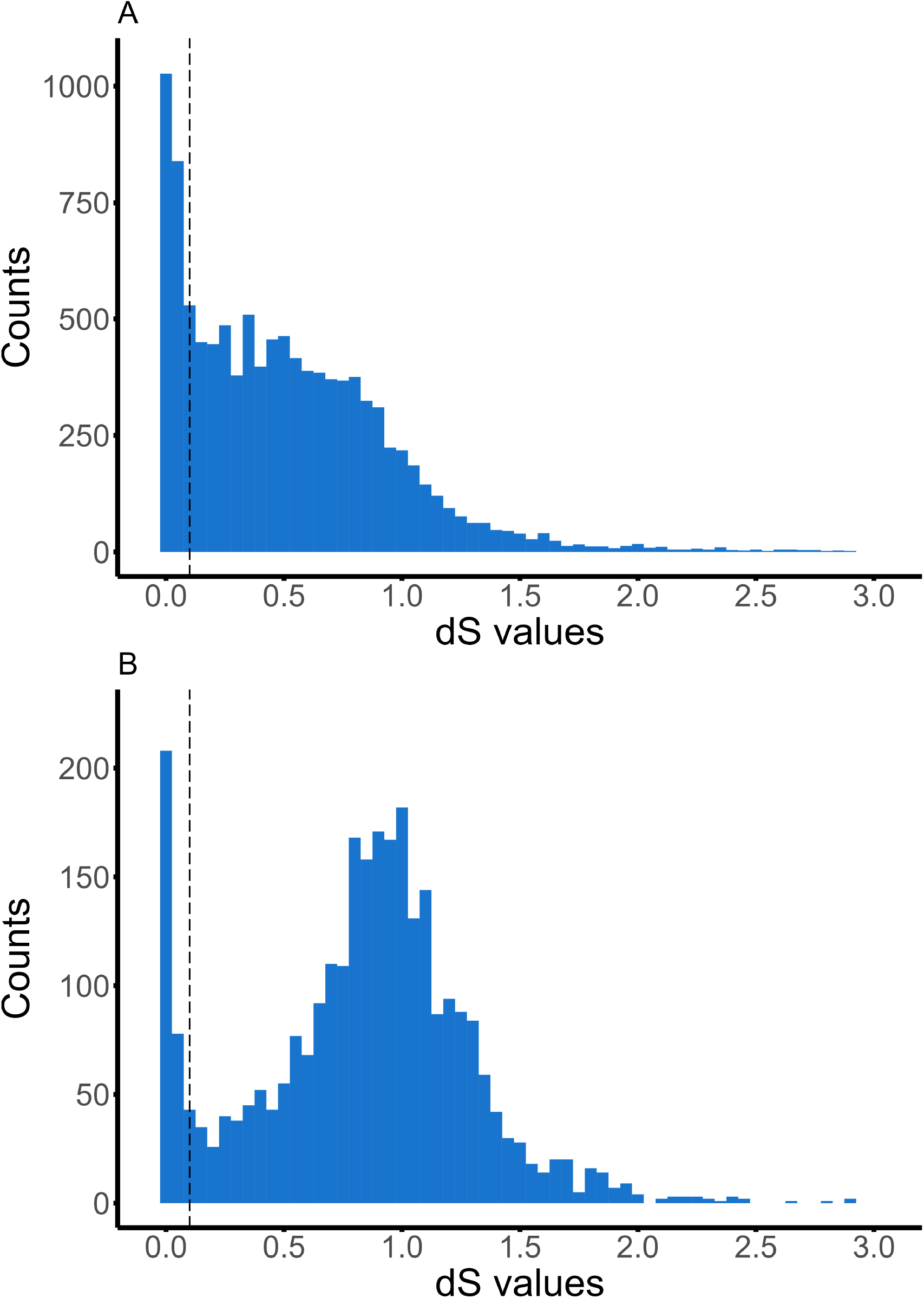
Distribution of *dS* of genes in cNUMTs. Shown are the distribution of *dS* between genes on the NUMTs and the host species mitochondrial DNA. NUMTs that possess at least one genes with *dS >* 0.1 are considered dcNUMTs, *dS* = 0.1 are indicated by a dashed line. Distributions are shown for **(A)** mammals and **(B)** birds. Note that count data for *dS* values >3 is not shown.

### Non-host origin of dcNUMTS reveal introgressed NUMTs

As recently integrated NUMTs will show little to no sequence divergence from their mitochondrial counterparts, they cannot tell us very much about the evolutionary consequences of NUMT integration. We hence focus on those cNUMTs that show substantial synonymous divergence to the respective gene in the host mitochondrial genome (dcNUMTs, divergent coding NUMTs with at least one gene with *dS >* 0.1 relative to the respective host mitochondrial gene), because these are likely the consequence of non-canonicial NUMT integrations (Figure 1). We identify 4,182 and 917 NUMTs that are dcNUMTs with 8,406 and 2,599 coding genes with *dS >* 0.1 in mammals and birds respectively, meaning that some dcNUMTs contain multiple divergent mitocoding genes.

We identify 148 dcNUMTs (3.54 %) in mammals and 249 dcNUMTs (27.15 %) in birds that we can assign to a non-host mitochondrial genome with high confidence (Table 1, Figure 4). Surprisingly we find that 77.19% and 40.46% of the dcNUMTs in mammals and birds, respectively, have no hit with any of the available mitochondrial genomes at the 95% identity level. To test whether these NUMT perhaps originate from species outside of the two taxonomic clades, we conducted further BLAST analyses but did not identify any further hits. The phylogenetic origins of these dcNUMTs are therefore located in their respective taxonomic group but the exact origins remain obscure. This suggests that these integrations are either not very recent, or that the current databases do not fully reflect the true biological mitochondrial diversity of mammals and birds.

**Figure 4:**
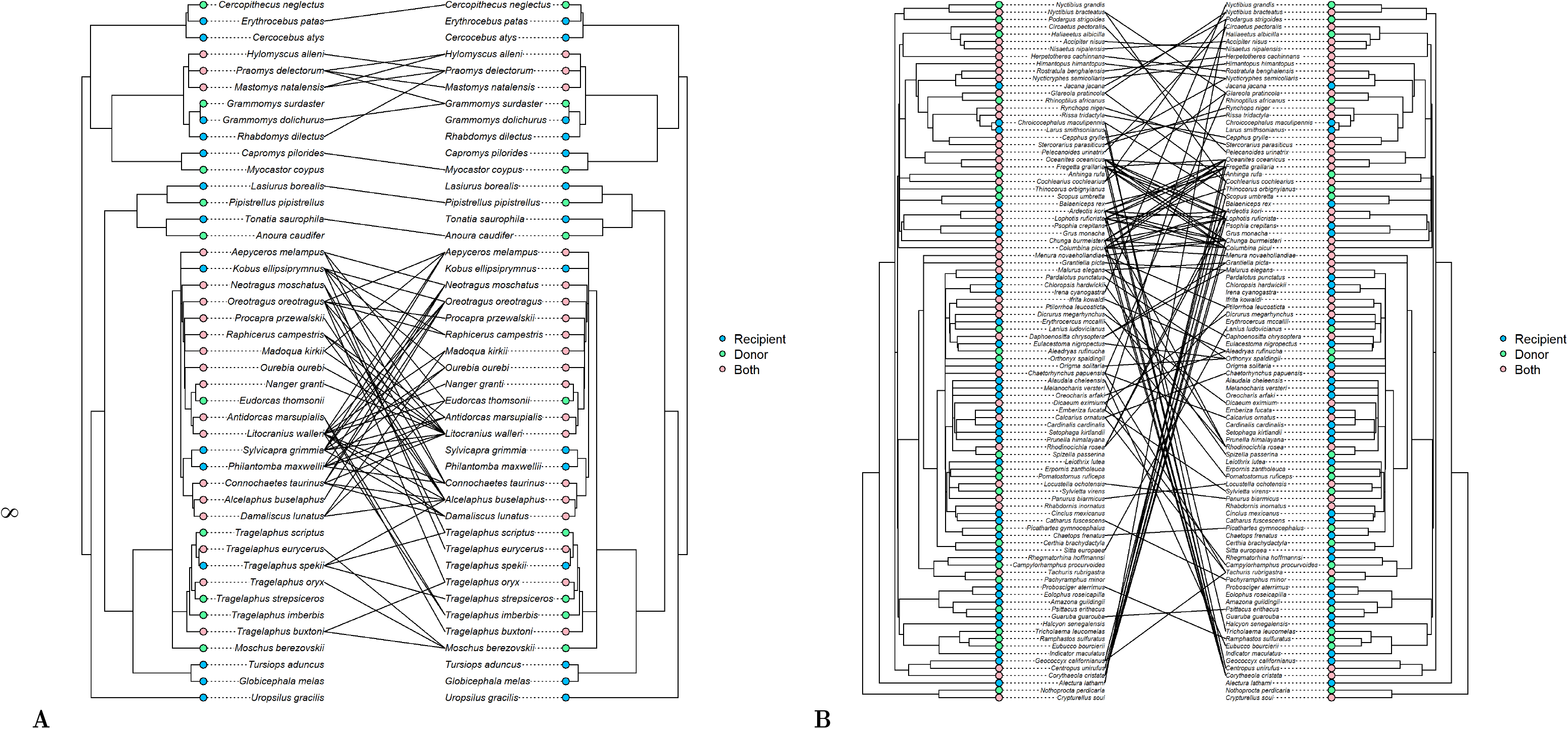
Phylogenetic pattern of NUMT integrations of non-host mitochondrial DNA. For **(A)** mammals and **(B)** birds. Shown are species that either show evidence of NUMTs of non-host origin or that appear be the donor species of the NUMT. Note that some species can be donor and recipient and that not all species included in our analysis are resolved phylogenetically in Timetree, so that not all pairs can be assigned. A complete list can be found in Tables S2 and S3.

**Table 1:** Numbers of BLAST hits of dcNUMTs against available mitochondrial genomes for birds and mammals. Best BLAST hit must have at least 95% identity to the query. *For a hit to be assigned as high confidence to a species other than the host, there needed to be a 5% or greater similarity to the non-host, e.g., a highly distinct hit to another species’ mitochondrial genome.

| Best Blast hit to (>95% identity, dcNUMT as query) | Mammals | Birds |
| --- | --- | --- |
| host species' mitochondrial genome | 441 | 18 |
| non-host species' mitochondrial genome | 513 | 528 |
| **with high confidence\*** | **148** | **249** |
| no species in the database | 3,228 | 371 |

#### dcNUMT gene analysis

Because dcNUMTS contain at least one gene with an intact reading frame, it is possible to investigate the evolutionary pressures that prevailed since the split from the last common ancestor using DNA codon models. For example, if genetic drift was the predominating force since the split from the last common ancestor of the NUMT and the host’ mitochondrial genome, one would expect the ratio of nonsynonymous to synonymous substitutions (*dN/dS*) to be approximately one. To test this, we divided our dataset into: (A) Single NUMT genes (Figure 5A), which highly divergent to other NUMT genes and therefore seem to be isolated (“single”); and (B) NUMT gene clusters (Figure 5B) that are highly similar, for instance due to shared speciation events (Vendrami et al. 2022). We then aligned the single NUMTs and NUMT gene cluster together with available mitochondrial genes of related species to calculate dcNUMT gene (Figure 5A) and gene cluster specific dN/dS ratios (Figure 5B) using codeml (Yang 2007). Codeml does not incorporate heterogeneity in codon composition or transition-to-transversion ratio (*κ*) which may differ between the nuclear and mitochondrial genome (Belle et al. 2005). However, we performed a series of simulations (Figure S1 and Table S1) that incorporate such heterogeneity (Fletcher and Yang 2009) which show little evidence for selection inference under a neutral model. We therefore consider our approach suitable to infer selection for nuclear integrated mitochondrial sequences.

**Figure 5:**
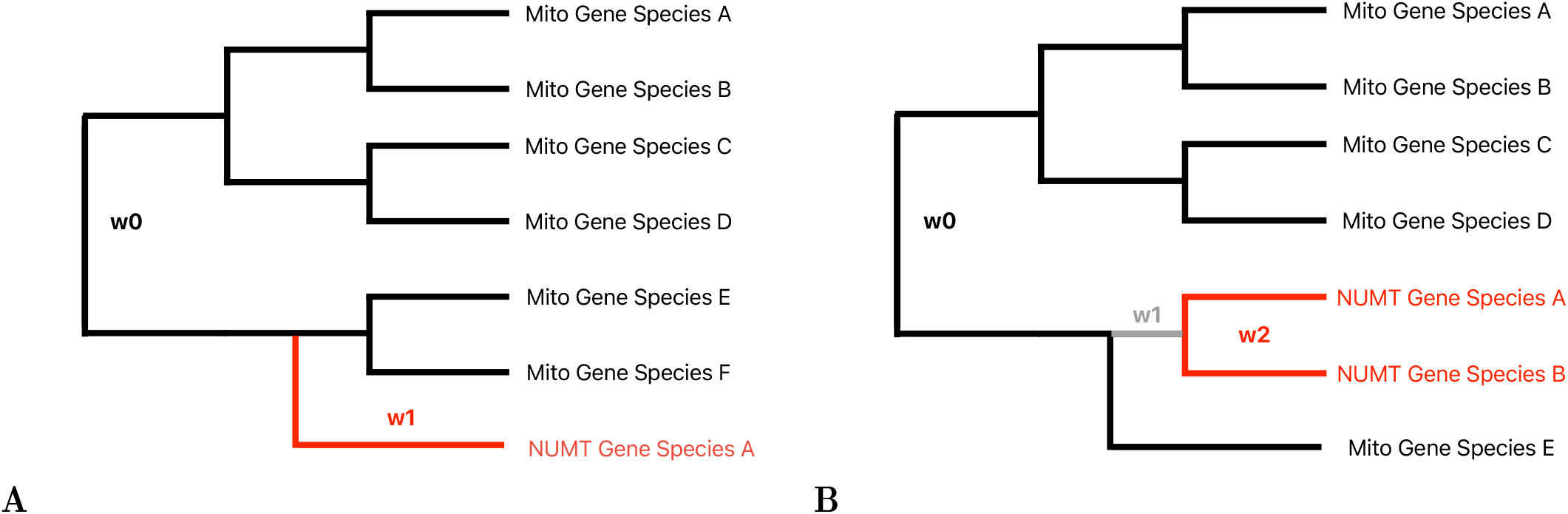
Illustration of the single gene and clustered genes codeml analysis. **(A)** Foreground branch in red (dN/dS=*ω*_1_) for a single NUMT gene, background branches (black, dN/dS=*ω*_0_) are for the respective mitochondrial genes of related species **(B)** Foreground branch in red for NUMT gene clusters (dN/dS=*ω*_2_, here illustrated with cluster size two), background branches (black, dN/dS=*ω*_0_ and grey, dN/dS=*ω*_1_) are for the respective mitochondrial genes of related species. Note that the grey branch is a seperate background branch, because it denotes the branch when the potential nuclear genomic integration took place.

#### Selection analysis on isolated dcNUMT genes suggest non-canonical NUMT integrations and “missing” mitochondrial genomes

We first conducted a codon analysis of the “single” dcNUMT genes. In mammals, we identified 3,549 single dcNUMT genes (42.2% of total dcNUMT genes) and 2,246 (86.4% of total dcNUMT genes) in birds. The differences in the proportion between the two taxonomic groups is likely because dcNUMTs genes in birds have greater synonymous divergence and are therefore generally more ancient (Figure 3). We conducted branch-specific analyses to test for positive and negative selection, as well as drift (Table 2). For the majority of single dcNUMT genes (Table 2), we identify sequence evolutionary patterns consistent with neutral evolution (62.4% for mammals and 83 % for birds). We also observe a smaller number of cases (37.6% for mammals and 17% for birds) that are consistent with purifying selection, and five cases of positive selection. However, it is possible that the last mitochondrial lineages from which these NUMT were derived are not included in our dataset, either because they became extinct or because they are lacking from our database. Therefore signatures of purifying selection for single NUMT genes could reflect the fact that our database lacks substantial parts of ancient or recent mitochondrial species diversity. However, our results are also consistent with the possibility of true purifying selection on the NUMTs.

**Table 2:** Evolutionary pressure analysis on single dcNUMT genes for mammals and birds. Shown are the numbers of genes that show branch-specific signatures of purifying selection, neutral evolution or positive selection.

| Evolutionary pressure on dcNUMT gene | # of genes in mammals | # of genes in birds |
| --- | --- | --- |
| Purifying selection ($dN/dS < 1$) | 1,337 | 377 |
| Neutral evolution ($dN/dS = 1$) | 2,211 | 1,865 |
| Positive selection ($dN/dS > 1$) | 1 | 4 |

#### Selection analysis on NUMT gene clusters suggest positive and negative selection on NUMTs

A proper evolutionary analysis of a single phylogenetically isolated NUMT gene is limited by the availability of the last mitochondrial lineage from which the mitochondrial DNA was transferred into a nuclear genome. To circumvent this limitation, we clustered the NUMT genes according to their similarity to each other independent of the host species. We then aligned these groups of gene sequences together with the most closely related mitochondrial genes available in our database. Based on these alignments, we then created gene trees and conducted clade-specific tests of selection (Table 3). We excluded gene trees that did not form monophyletic NUMT clusters. We then tested the selection pressure acting on the clade representing the diversification of the NUMTs (i.e. a node based last common ancestor approach (Lee 1998), in contrast to branch-based last common ancestor, e.g. excluding the grey branch in Figure 5B). By that it is likely that the considered clade reflects an evolutionary time when the NUMT was truly integrated into the nuclear genome, e.g. a pattern one would observe due to speciation events (Vendrami et al. 2022). Consequently would the estimated selective pressure analysis reflect selection on the NUMT, but not mitochondrial DNA.

**Table 3:** Evolutionary pressure analysis on dcNUMT gene clusters for mammals and birds. Shown are the numbers of gene clusters that show clade-specific signatures of purifying selection, neutral evolution and positive selection, respectively.

| Evol. pressure on dcNUMT gene clusters | # of clusters mammals | # of clusters in birds |
| --- | --- | --- |
| Purifying selection ($dN/dS < 1$) | 247 | 24 |
| Neutral evolution ($dN/dS = 1$) | 726 | 73 |
| Positive selection ($dN/dS > 1$) | 23 | 1 |

We identified over ten times more of these NUMT gene clusters for mammalian species than for birds (996 versus 98). The average number of species with NUMT genes per cluster was three for both taxonomic groups (2.98 and 3.01, for mammals and birds respectively). While mammalian dcNUMTs are generally less divergent to the host’s mitogenome than avian dcNUMTs (Figure 3), they are as frequently shared between species. Based on the evolutionary rate analysis we identify more than 250 gene clusters that show evidence of purifying selection, most of them specific to mammals. Moreover, we identify an additional 24 gene clusters, 23 for mammals and 1 for birds, that show evolutionary signatures consistent with positive selection. This suggests that the identified NUMTs are potentially functionally conserved and therefore contribute towards adaptive variation.

#### Evidence of positive selection in a human NUMT

We identified one NUMT on chromosome 2 in humans that is part of a ND4L gene cluster with evidence for positive selection (Table 4, one out of 23 NUMT clusters for mammals). This particular NUMT is shared among seven ape species (*Nomascus leucogenys*, *Pan troglodytes*, *Gorilla gorilla gorilla*, *Pan paniscus*, *Homo sapiens*, *Hylobates moloch*, *Pongo abelii*) but the NUMT itself clusters with mitochondrial sequences of species of the family Callitrichidae (Figure 6). In the human genome, this NUMT is flanked by genes encoding for ARHGAP15, GTDC1, ZEB2, KYNU and LRP1B. We also conducted an alphafold structural prediction (Mirdita et al. 2022) using the NUMT’s gene sequence translated into a protein with the vertebrate mitochondrial code (Figure S2) and find very little 3D structural variation between the human ND4L protein and the 3D protein prediction of the NUMT gene. Therefore, this hypothetical prediction does not suggest that the accumulated mutations in the NUMT strongly alters the three-dimensional structure. It therefore appears unlikely that the signal of positive selection of the mitocoding nuclear DNA is the consequence of a false positive, e.g. through misalignment or rapid accumulation of random amino-acid altering mutations (Mallick et al. 2009). Instead we find that the genomic region where the NUMT is located is known to have evolved under positive selection in the ancestral lineage of modern humans (Green et al. 2010; Prüfer et al. 2013; Racimo 2015).

**Figure 6:**
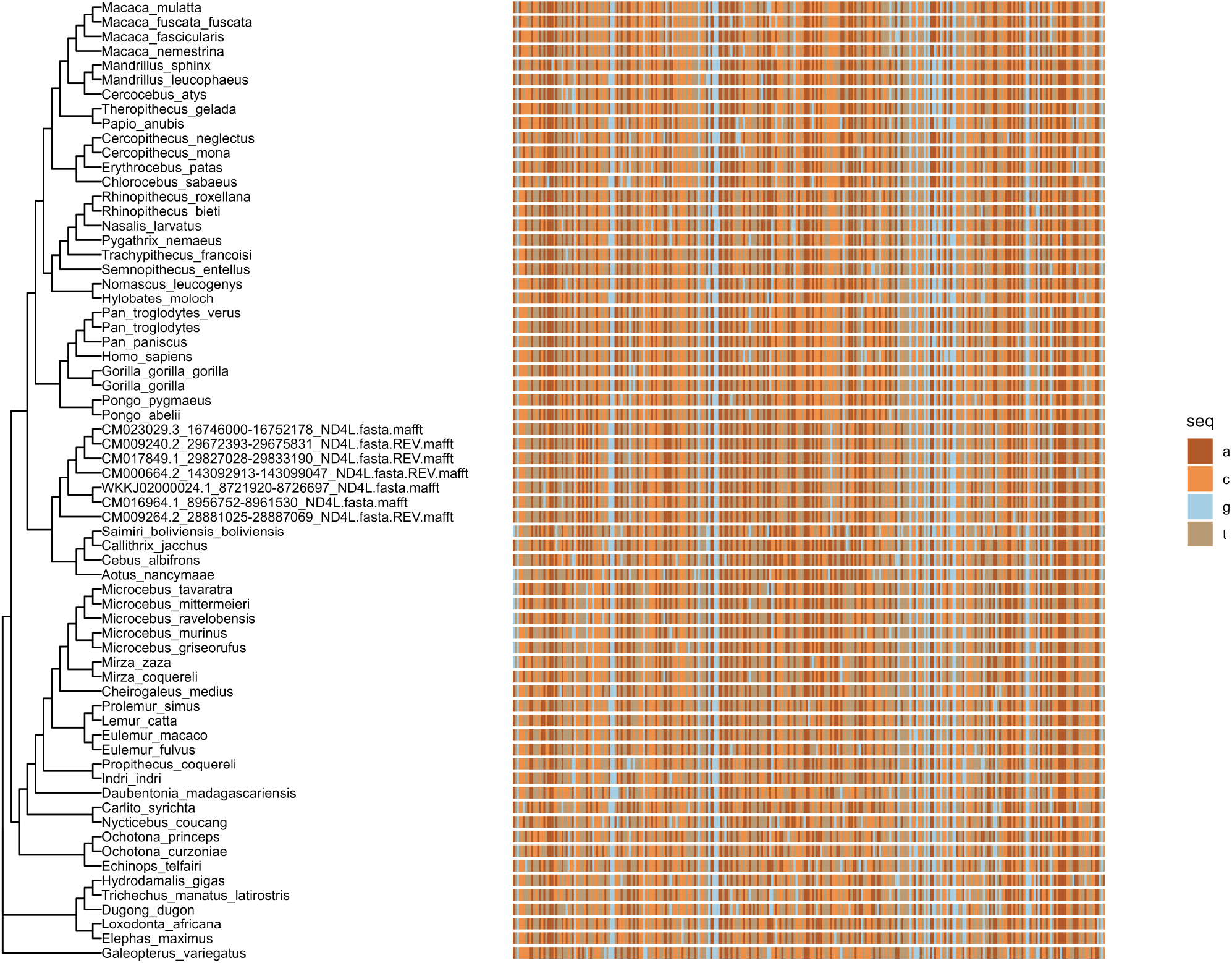
Phylogenetic tree and sequences used for codeml analysis of a NUMT cluster that contains dcNUMTs of seven ape species along with mitochondrial sequences of 59 (sub)species. The NUMT gene with an intact reading frame is ND4L. Evolutionary pressures were estimated with codeml using the mitochondrial genetic code. The coding sequence is shown in color using the 4-letter-DNA notation illustrating no pinpointed divergence of the NUMT sequences.

**Table 4:** Codeml analysis results of a mitocoding gene located on a NUMT that is shared among seven ape species. Rate heterogeneity between the NUMT clade and the mitochondrial sequences was inferred by comparing a two-ratio model with a one-ratio model (single dN/dS=*ω* value for the entire tree). Positive selection was inferred from a two ratio model and a fixed two-ratio model for which the *dN/dS* of the NUMT subtree was fixed to *ω*_2_=1. The tree and sequences used in the analysis are shown in Figure 6.

| Model | ln Likelihood | $\omega$ values | Comparison |
| --- | --- | --- | --- |
| One-ratio | $\ln L_1=6,946.87$ | $\omega = 0.06$ | |
| Two-ratio (fixed) | $\ln L_{2a}=6,905.41$ | $\omega_1 = 0.05, \omega_2 = 1$ | |
| Two-ratio | $\ln L_{2b}=-6,902.98$ | $\omega_1 = 0.05, \omega_2 = 2.9$ | $2\Delta(\ln L_{2a}, \ln L_{2b})$<br>$P=0.027 (\chi^2, d.f.=1)$ |

## Discussion

Here, we present a comprehensive analysis of mitochondrial DNA that has been integrated into the nuclear genomes (NUMTs) of more than 1,000 mammalian and avian species. We use a previously established pipeline that uses a species’ mitochondrial genome as a query for a nuclear genomic BLAST identification of the same species (Vendrami et al. 2022). Altogether we identify more than 80,000 NUMTs in mammals and more than 29,000 NUMTs in birds (Table 5). Generally, there are more NUMTs in mammalian genomes, which is not necessarily surprising given that mammalian genomes tend to be larger (Kapusta, Suh, and Feschotte 2017). We also find a positive relationship between genome size and the number of mitochondrial integrations in mammals but not birds. A possible explanation for this is that mammals contain NUMT copies resulting from intragenomic duplications. Birds are known to exhibit very little variation in their genome sizes in general, which might also explain the lack of intragenomic copies, such as segmental duplications. For many of the identified NUMTs, it is unclear on which chromosome in the genome they are located, as they mainly located on small scaffolds that are often not much bigger than a mitochondrial genome (e.g. <20kb) itself (Figure S3). However, large-scale application of long-range haplotype-based genomic sequencing (Rhoads and Au 2015; Lu, Giordano, and Ning 2016) will overcome the current limitations of identifying the precise genomic locations of nuclear DNA of organelle origin in the near future.

**Table 5:** Number of NUMTS, coding NUMTs (cNUMTs) and deep coding NUMTs (dcNUMTS) in birds and mammals. As dcNUMTs can have multiple genes they might group with other dcNUMTs multiple times.

| NUMT type | Number in mammals | Number in birds |
| --- | --- | --- |
| NUMTS | 85,100 | 28,947 |
| cNUMTS | 5,229 | 1,200 |
| dcNUMTS | 4,182 | 917 |
| dcNUMTS/non-dcNUMTS in groups | 2,670/144 | 157/6 |
| single dcNUMTS | 2,024 | 786 |
| dcNUMTS single+grouped | 511 | 26 |

Some of the identified NUMTs are very distantly related to the host’s mitochondrial DNA. Because our approach of identifying NUMTs tends to detect NUMTs that show a reasonable degree of similarity between the NUMT and the host’s mitochondrial genome, it is likely that we did not identify all distantly related NUMTs. Therefore, our estimates of the NUMTs per species are potentially underestimates. For example, we do not find any NUMTs that were highly similar to a species outside of the respective animal class. Because we do not attempt to identify NUMT integration events as such, some of the NUMTs may result from structural mutations, such as segmental genome copies. Hence our NUMT numbers are an overestimate of the number of NUMT integration events from non-nuclear DNA. However, because we focus on dcNUMTs for our evolutionary analysis it is irrelevant whether a NUMT has undergone intragenomic copying or is the result of a “original” genomic integration.

Our method also depends on an accurate database assignment of the mitochondrial genome to the respective species. However, due to biological mitochondrial diversity (Clark et al. 2023) or simply erroneous entries in databases (Caswell et al. 2019; Sangster and Luksenburg 2021), mitochondrial DNA and the NUMT DNA could appear more divergent. To limit the impact of this to our dcNUMT analysis, we focused on coding NUMTs that have at least 10% synonymous divergence to the respective mitochondrial gene. To investigate the possibility of erroneous database entries we checked the mitochondrial genomes of 22 bird species that show a high number of highly dissimilar NUMTs. For this, we assembled the mitochondrial genomes of these species from raw sequencing data using an established pipeline (Jin et al. 2020). While we observed some genetic variation between the assembled and the canonical mitochondrial genome in the database (Table S4), we did not find any evidence for an incorrect assignment of mitochondrial genomes. We conclude that falsely assigned mitochondrial genomes play a very limited role in our analysis.

### dcNUMTs

To identify non-canonical origins of NUMTs and potential signatures of selection, we focused on dcNUMTs, which still contain genes with an intact mitochondrial reading frame. Because the nuclear and mitochondrial genome use different genetic codes, it is unlikely that mitochondrial genes integrated into the nuclear genome can be translated into intact proteins. Moreover, after integration of the mitochondrial DNA into the nuclear genome, NUMTs are expected to be accumulate mutations over time that likely disrupt the mitochondrial reading frame. However, we find more than 5,000 NUMTs in mammals and birds (Table 5) that contain genes with an intact mitochondrial reading frame and that show relatively high divergence to the respective hosts mitochondrial encoding genes (dcNUMTs with genes with *dS >* 0.1, i.e. a synonymous per site divergence of more than 10%). For mammals we find that the number of dcNUMT becomes lower for increasing dS values but in birds there is a clear peak at dS≈ 1. This might reflect a historic episode of high transposable element activity common to all birds. However, as the nuclear and mitochondrial mutation rates may vary substantially across species, the *dS* estimates should be considered only as a rough proxy of the integration time of the NUMT.

It is possible that dcNUMTs are transcribed and may have a function on the RNA level, or that the DNA itself has some regulatory function as observed for mitochondrial DNA (Acker et al. 2023) or creates novel exons (Noutsos et al. 2007). For example, the transcription of non-coding genomic DNA is very common (Gao et al. 2020) and active retroviruses (Bolisetty et al. 2012) may could lead to expression of NUMTs as well. However, this remains speculative and would need to be supported by a signature signature consistent with functionality. To address this, we investigated the evolutionary signatures of mitocoding NUMT genes. Specifically, two separate analyses that may explain the existence of dcNUMTs were conducted. First, we focused on the hypothesis that the NUMTs result from introgression or hybridisation events. Second, we analysed the selection pressures on coding DNA to identify traces of directional selection.

### Introgression/Hybridisation

To identify potentially introgressed/hybridised NUMTs we BLASTed the dcNUMTs against available mitogenomes, that also contain mitochondrial genomes that were not in our initial analysis (e.g. because there was no nuclear genome available for the species). We identify 67 dcNUMTs in mammals and 132 in birds that can be assigned to a different other species than the host’s species mitogenome with high confidence (Tables 1, S2 and S3 as well as Figures 4A and 4B). We observe a greater number of distant hits in birds than mammals. This is possibly due to the fact that the bird genomes in our dataset diverged earlier than the mammal genomes - which means that their mitogenomes are more diverged, allowing for better discrimination of non-canonical NUMT origins. In mammals, almost all of the introgressed NUMTs occur within the ungulates (Figure 4A and Table S2) - a taxonomic group in which hybridization plays an extensive role (Iacolina et al. 2018).

In birds, we find introgression events between distantly related species (Figure 4B, Table S3). For example, we identified a NUMT that shows a signature of introgression from the tawny frogmouth (*Podargus strigoides*), a species of frogmouth native to the Australian mainland and Tasmania, to the gweela (*Alectura lathami*), a mound-building bird from the family Megapodiidae found in eastern Australia. A possible explanation of cross-species NUMT transfer might be that some NUMTs co-localise with retrotransposon elements (Tsuji et al. 2012; Bolisetty et al. 2012; Dayama et al. 2014; Schiavo et al. 2017), and the cross species transmission of retroviruses has been documented in mammals and non-vertebrates (Flavell 1999; Diehl et al. 2016). We also note that the W chromosome is a refugium for retroviruses in birds (Peona et al. 2021), which might imply that we may have missed NUMTs that are located on the W if the respective genome of a species was obtained from a male individual, which has the homogametic genome in birds. Because our dataset heavily relies on genome assemblies retrieved from short-read sequencing, we cannot pinpoint the exact genomic locations of many of the identified NUMTs, because they tend to be located on very short scaffold (Figure S3) - and this appears to particularly true for avian dcNUMTs. Unable to locate the precise genomic locations, however, hampers the possibility to pinpoint integration events that happened as a consequence of genomic hybridization. In the future this can be overcome by the application of long-range sequencing approaches, which will subsequently allow us to use dcNUMTs as markers for genomic hybridisation and could help to further unravel their potential role as adaptive elements.

We also identify 3,179 and 384 dcNUMTs in mammals and birds, respectively for which we cannot identify a homologous sequences with a sequence identity of more than 95% (Table 1). While this could mean that these are simply older NUMTs that are highly diverged but which have remained in the genome for a long time, it could also mean that some mitogenomes are lacking from our database. This is either because the respective species has not been yet sampled or because the respective species has already gone extinct. Surprisingly, the number is almost ten-fold higher in mammals compared to birds. A deeper mammalian sampling or the retrieval of ancient mammalian mitogenomes might be enable us to reveal the evolutionary origins and fates of further mammalian NUMTs in the future.

### Tests of selection

To test for potential signatures of selection specific to NUMTs we conducted substitution rate analysis of genes located in NUMTs. We restricted our analysis to those genes with intact reading frames showing substantial synonymous divergence to the host’s mitochondrial genome. Specifically, we conducted our analysis using the mitochondrial genetic code as there is no known mechanism that would allow nuclear genes to be translated into proteins using the mitochondrial code. Instead, it is likely that the reading frames of NUMT genes should deteriorate over evolutionary time due to the acquisition of new mutations. The long-term maintenance of NUMT genes might therefore be a signal of selection. For this, we classified the NUMT genes into two groups, single NUMT genes (i.e. isolated NUMT genes with no other NUMT genes in close proximity) and grouped NUMT genes (i.e. NUMT genes with high similarity to a NUMT gene of another species or another NUMT gene of the same species). The latter group would allow us to evaluate the evolutionary pressures that have acted on the NUMT gene when it integrated into the nuclear genome (Figure 5).

For the majority of single NUMT genes, we observe a signature of selection consistent with drift (e.g. dN/dS=1). However, we also identify more than 2,000 genes showing signatures of purifying selection (Table 2), comprising 54% and 83% of the tested NUMT genes for mammals and birds, respectively. Surprisingly, we also identify five NUMT genes showing signatures of positive selection. While it is intriguing that many genes have been evolving under selection pressure, this analysis is hampered by the fact that for the tested branches, the single NUMT genes may have existed as mitochondrial genome sequences. Unfortunately we cannot incorporate the precise date of the nuclear integration, as we potentially lack a substantial amount of mitodiversity (Table 1) and therefore cannot be sure that we include a sequence that did not divergence before the NUMT integration took place. To address this limitation, we conducted clade specific tests of selection on groups of NUMT genes. By that it is likely that the considered branches in the clade with the NUMTs reflect an evolutionary time when the NUMT was truly integrated into the nuclear genome. However, it is also possible that by that approach we group sequences that have evolved the same sequence independently (Hazkani-Covo, Zeller, and Martin 2010; Song et al. 2013). Again, for the majority of grouped NUMT genes we find an evolutionary signature consistent with genetic drift (Table 3), with 64% and 68% of the groups showing selection pressures consistent with drift for mammals and birds, respectively.

### Positive selection in a human NUMT gene

Finally, we identified one gene within a human NUMT that is part of a cluster of seven ape NUMTs. For humans, the genes located in the genomic region where the NUMT is located are known to have evolved under positive selection in the ancestral lineage of modern humans (Green et al. 2010; Prüfer et al. 2013; Racimo 2015). Moreover, structural mutations, such as deletions and duplications, in the respective region are known to cause the Mowat-Wilson syndrome in humans (Baxter et al. 2017; Goyal et al. 2022), a rare but severe genetic disorder. Because the NUMT is linked to functionally important genes, it is a possible explanation of the signal of positive selection - the NUMT might be hitchhiking with a gene under positive selection (Barton 2000). Hence, the fact that one mitochondrial gene on the NUMT shows a signature of positive selection at the (mitochondrial) protein coding level that is shared between many ape species, fuels speculation to a potential functional role of this NUMT or one of its genes.

## Conclusion

Here we have shown that NUMTs may originate from non-canonical introgression and that signatures of purifying and positive selection are common and widespread across two main vertebrate clades. Currently, in many genome assemblies NUMTs cannot be placed onto chromosomes, because they are usually flanked by highly repetitive sequences due to retro-virus integration. Haplotype based sequencing will likely be able to place more NUMTs on chromosomal locations, and thereby reveal the evolutionary signatures of NUMTs as a consequence of linkage. Our example of a human NUMT has shown that it is crucial to know the genomic context to deduce the likely functional roles of these intriguing genetic elements. Moreover, recent and ancient introgression maybe understood better when placing NUMTs into phylogenetic context. Even if the NUMTs themselves are not functional, they provide excellent markers of (positive) selection and introgression/hybridisation.

## Materials and methods

### Data collection and selection

The genome assemblies of all mammalian species (available before November, 11, 2021) species and all avian species (available before November, 02, 2021) species were obtained from GenBank (Sayers et al. 2021) using ncbi-genome-download v0.3.1 (https://github.com/kblin/ncbi-genome-download). If one species had multiple assemblies, the representative genome or the genome with higher coverage (if no assemblies were stated as representative) was included in our database.

The mitogenomes with at least one partial coding gene of all mammalian species (available before November, 22, 2021) or all avian species (available before December, 16, 2021) were obtained from GenBank for building BLAST databases. The mitogenomes from the same species and subspecies (or closely related subspecies if the same subspecies were not found) with genome assemblies were selected for NUMT identification. If one species had multiple mitogenomes, the reference assembly was included. Unverified mitochondrial genomes were ignored. The protein-coding gene annotations were also obtained from GenBank. The mitogenomes without protein-coding gene annotations were annotated using the MITOS2 webserver (Donath et al. 2019). See Table S5 for the complete list of genomes and mitochondrial genomes and their accession numbers.

### NUMT identification

For NUMT identification, we performed local BLASTN v2.6.0+ (Camacho et al. 2009) with the word size of 20 and used mitochondrial genomes as queries to search the corresponding genome assemblies of the same species. This is a modified approach previously published(Lammers et al. 2017; Vendrami et al. 2022). The hits with subject lengths shorter than 200 bp were discarded. The corresponding NUMT sequences were extracted with BEDTools v2.26.0 (Quinlan and Hall 2010). NUMTs and their reverse sequences (generated by EMBOSS v6.6.0.0 revseq algorithm (Rice, Longden, and Bleasby 2000) were pairwisely aligned with the protein coding genes of the corresponding mitochondrial genome using MAFFT v7.310 (Katoh and Standley 2013) to identify cNUMTs.

The NUMTs numbers were mapped to the species trees from Timetree (Kumar et al. 2017) to examined their evolutionary trends across the phylogenies. The rate of evolution were estimated with independent contrast model using reversible jump Markov chain Monte Carlo (RJMCMC) in BayesTraits v.4.0.1 (Pagel and Meade 2022). For each dataset we ran 12,000,000 iterations, sampling once every 2,000 iterations after a 2,000,000 burn-in iterations. The branch length transformation based on the rate change were applied to ancestral reconstructions of NUMTs numbers with ace function in the phytools packages (Revell 2012) in R (v4.2.3).

### Substitution rate analyses of NUMTs with coding genes

We first checked whether the cNUMTs in pairwise alignment with the corresponded mitochondrial protein coding genes were coding in frame (e.g. lacked internal stop codons). The inframe cNUMTs and their corresponded mitochondrial protein coding genes were used to calculate the synonymous (dS) substitution rates with KaKs calculator (Wang et al. 2010) under the NG substitution model (Nei and Gojobori 1986). The NUMTs with at least one coding gene with dS ≥ 0.1 were considered as potentially dcNUMTs. These potentially dcNUMTs were used as queries to BLAST against (Camacho et al. 2009) the mitogenomes of all mammalian species and all avian species, respectively (Table S5). The best Blast hits were define as a Blast hit with the highest bit-score. The high confidence best Blast hits were define as a best Blast hits with at least 95% identity to the query. The dcNUMTs with high confidence best Blast hits to mitogenomes from another species with the differences of percentages of identical matches ≥ 5 to the Blast hits to their host species were considered potential lateral transfer between species pairs.

### Clustering NUMTs and selection analysis

To investigate the functional constraints of coding sequences on cNUMTs, we first identified NUMTs derived from the same source and then analyzed the ratio of non-synonymous to synonymous (dN/dS) of each NUMT clusters. We used mafft v7.310 (Katoh and Standley 2013) to align protein coding sequences identified in cNUMTs and reconstructed the coding sequence trees with iqtree v2.0.3 (Minh et al. 2020). Then we used CODEML implemented in PAML (Yang 2007) to calculate pairwise dN/dS comparisons and substitutions per codon on all the protein coding genes from cNUMTs which corresponding to the same mitochondrial protein coding genes. The coding sequence pairs with substitutions per codon less than 0.1 were examined for their best hits against the mitogenomes. The pairs with both coding sequences’ percentage of identical matches less than 98% were examined if any of the coding sequences matched with coding sequences in other pairs. The coding sequences matched in pairs were grouped into one cluster.

We then translated the mitochondrial protein-coding genes from the species we used to identify NUMTs with Biopython (Cock et al. 2009) and aligned each mitochondrial protein-coding genes with mafft v7.310 (Katoh and Standley 2013). The translated seqeunces with length 20% shorter than the longest sequnces in the alignment were removed to exclude incomplete mitochondrial protein sequences. The alignments of corresponded mitochondrial protein-coding genes were used to align with each dcNUMT clusters generated in the previous step with prank v.170427 (Löytynoja 2014). The alignments were used to reconstruct coding sequence trees with FastTree v2.1.11 (Price, Dehal, and Arkin 2010). In order to increase computational efficiency, we extracted the subtrees with the clades containing cNUMT genes and 3 hierarchical levels above the focal clades using Newick Utilities v1.7.0 (Junier and Zdobnov 2010). The codon alignments of the corresponding subtrees were converted from mitochondrial gene sequences and their translated protein sequences using pal2nal v14 (Suyama, Torrents, and Bork 2006). Then we used CODEML implemented in PAML (Yang 2007) for the selection analysis. We implemented two different branch models: a three-ratio model (model = 2, NSsites = 0) and a three-ratio model with the omega of NUMT clades fixed to 1 (model = 2, NSsites = 0, fix_omega = 1, omega=1), for each cNUMT gene cluster. The likelihood differences among the models were used to derive statistical significance. Benjamin-Hochberg adjustments (Benjamini and Hochberg 1995) were applied for multiple testing corrections.

We also conducted a set of analysis to test. Potential differences in mitochondrial and nuclear DNA might be reflected in the transition to transversion ratio (ts/tv, *κ*) which is much higher in mitochondrial DNA relative to nuclear DNA in humans, although there is lots of variation across animals (Belle et al. 2005). While codeml does not incorporate heterogeneity in *κ* or codon composition, it is possible to conduct sequence simulation using branch specific models of sequence evolution (Fletcher and Yang 2009). For this we performed simulations with INDELible and used estimated *κ* and codon frequencies from our data (i.e. cluster depicted in Figure 6, with NUMT and mitochondrial specific codon composition and *κ* estimated separately) as inputs. We also simulated a homogenous model where we assumed codon composition and *κ* from the mitochondrial sequences only. We then tested these simulated sequences using codeml that assumes homogeneous codon composition and a single *κ* across branches similar to our test setup (Figure S1 and Table S1).

## Data deposition

Scripts are available at github https://github.com/chnyuch/numt_vertebrate and https://github.com/tgossmann/numt_vertebrate.

## Acknowledgment

This project has received funding from the European Research Council (ERC) under the European Union’s Horizon 2020 research and innovation programme grant agreement No. 947636. This work was supported by the BMBF-funded de.NBI Cloud within the German Network for Bioinformatics Infrastructure (de.NBI) (031A532B, 031A533A, 031A533B, 031A534A, 031A535A, 031A537A, 031A537B, 031A537C, 031A537D, 031A538A).

## Author contribution

TIG designed the study with input from JIH and DV. YC conducted the majority of the analyses. MH and LH conducted analyses on species subsets. TIG and YC drafted the manuscript with input from all of the authors.

## Appendix Supplementary Figures

**Figure S1:**
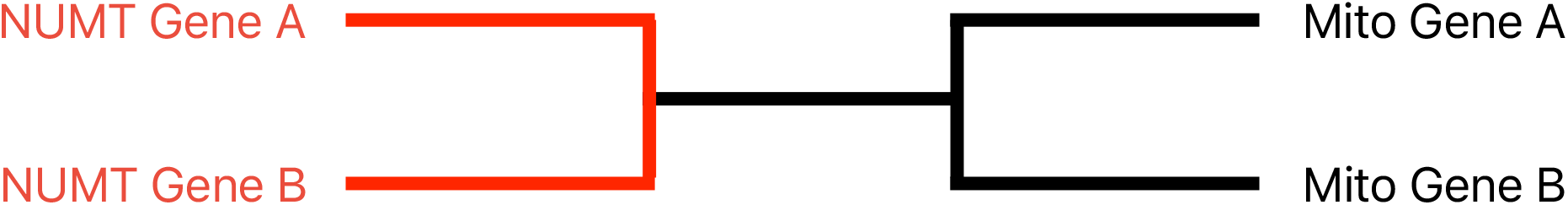
Tree used to conduct heterogenous sequence simulations with INDELible. Sequences were simulated under the base composition derived from the ND4L gene cluster that contain mitochondrial as well as mitonuclear gene sequences (Figure 6). Base composition and transition-transversion ratio (*κ*) were seperately estimated for mitochondrial sequences and NUMT sequences. These estimates were then used to simulate a heterogenous model where red branches were evolved under the estimates for the NUMT sequences and the black branches under the estimates derived from the mitochondrial sequences. *dN/dS* was varied for each simulation and was tested for non-neutral evolution using the codeml setup used in this study.

**Figure S2:**
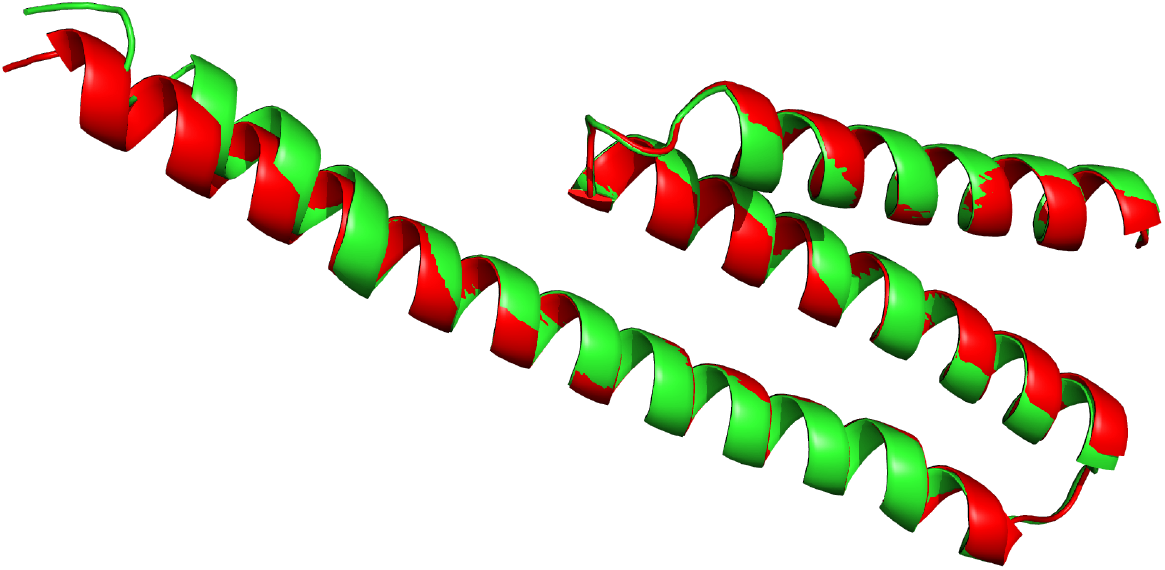
Alphafold 3D prediction of a human NUMT gene along with the corresponding human mitochondrial gene. The NUMT gene is part of a dcNUMT cluster that is shared among seven ape species (Figure 6). The NUMT gene with an intact reading frame is ND4L. Shown are the 3D structures of human ND4L as well as a structure predcition conducted with alphafold based on the hypothetical protein seqeunce assuming mitocondrial code

**Figure S3:**
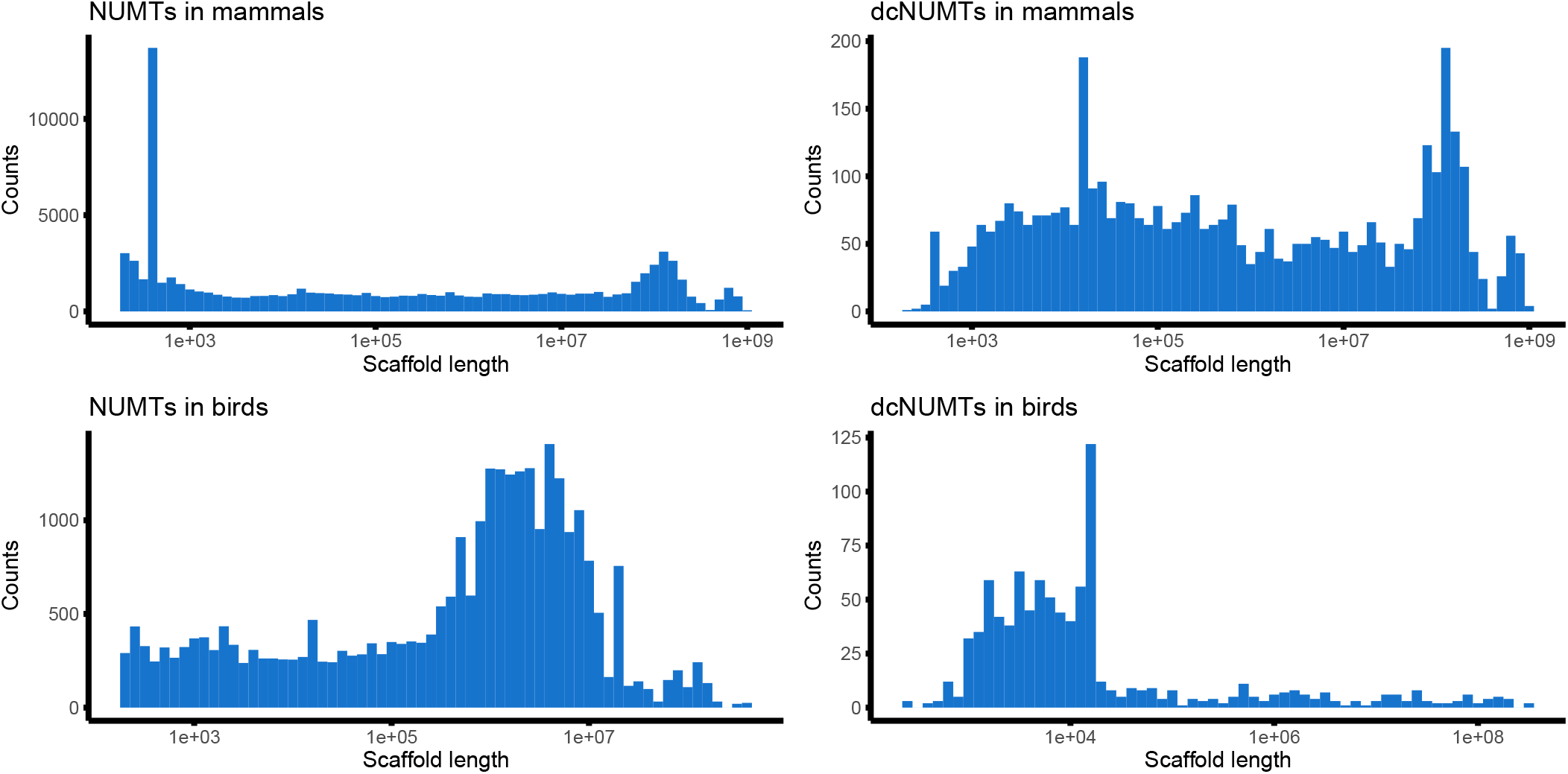
Distribution of genomic scaffold length of scaffolds that contain NUMTs. shown for mammalian and avian species. Many identified NUMTs are on short genomic fragments that are not assigned to chromosomes.

## Supplementary Tables

**Table S1:** Codeml model comparison of sequences simulated under heterogeneous codon composition and transition-transversion (κ) ratio as well as homogeneous compositions. Sequences were simulated under the tree depicted in Figure S1 using INDELIBLE. *ω* ratios were inferred and tested for significant differences using a two-ratio model with a set of foreground and background branches S1. *ω* ratio for the background ratio was constant (*ω_B_* = 0.04, estimated from the ND4L cluster depicted in Figure 6). For the homogeneous model codon composition and *κ* was simulated under the estimates derived from mitochondrial sequences.

| foreground branch | heterogeneous | homogeneous |
| --- | --- | --- |
| simulated $\omega_F$ | significant ($\omega_F \neq 1$) | significant ($\omega_F \neq 1$) |
| 1 | 0% | 0% |
| 0.8 | 7% | 1% |
| 0.6 | 57% | 68% |
| 0.4 | 98% | 97% |
| 2.25 | 96% | 94% |

**Table S2:** All mammalian host - donor pairs of NUMTs that likely originated from a different species. For full table see Supplementary_tables.xlsx

| Recipient species | NUMT ID | Donor scaffold ID | Donor species |
| --- | --- | --- | --- |
| Aepyceros melampus | SJXQ01006944.1:63574-66381 | NC_020699.1 | Connochaetes taurinus |
| Aepyceros melampus | SJXQ01006944.1:78349-91308 | NC_039669.1 | Eudorcas thomsonii |
| Aepyceros melampus | SJXQ01012544.1:217760-229605 | NC_020728.1 | Neotragus moschatus |
| Alcelaphus buselaphus | SJXN01419338.1:1-924 | NC_020731.1 | Oreotragus oreotragus |
| Alcelaphus buselaphus | SJXN01502202.1:1-1258 | NC_020731.1 | Oreotragus oreotragus |
| Alcelaphus buselaphus | SJXN01519847.1:1-1458 | NC_020731.1 | Oreotragus oreotragus |

**Table S3:** All avian host - donor pairs of NUMTs that likely originated from a different species. For full table see Supplementary_tables.xlsx

| Recipient species | NUMT ID | Donor scaffold ID | Donor species |
| --- | --- | --- | --- |
| Accipiter gentilis | BJER01014652.1:2587-13711 | NC_040858.1 | Haliaeetus albicilla |
| Accipiter nisus | BJBX01032538.1:1946-9915 | NC_007598.1 | Nisaetus nipalensis |
| Alaudala cheleensis | VWYE01023216.1:285-2530 | MN356176.1 | Eubucco bourcierii |
| Alaudala cheleensis | VWYE01025130.1:1181-2744 | MN356176.1 | Eubucco bourcierii |
| Alectura lathami | VXAV01019081.1:1-243 | MN356205.1 | Podargus strigoides |
| Amazona guildingii | VXAR01004813.1:3-2347 | MN356178.1 | Ciccaba nigrolineata |

**Table S4:**
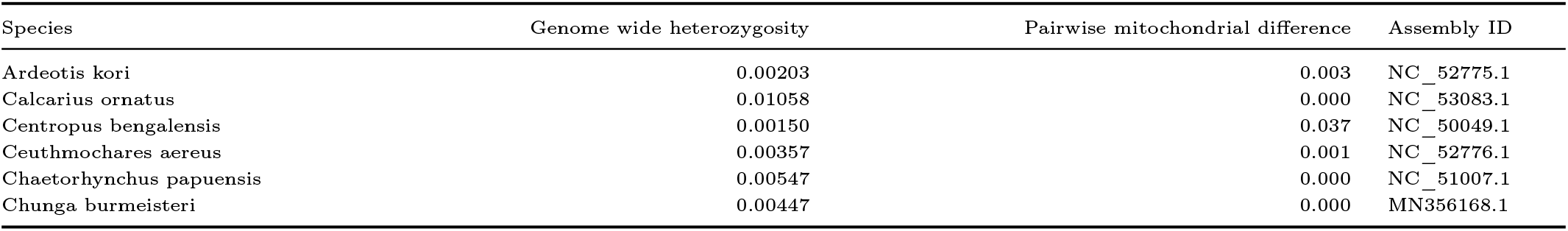
Comparison of the genetic and mitogenetic diversity of 22 avian species for which mitogenomic diversity was retrieved form raw sequencing reads. For full table see Supplementary_tables.xlsx

**Table S5:**
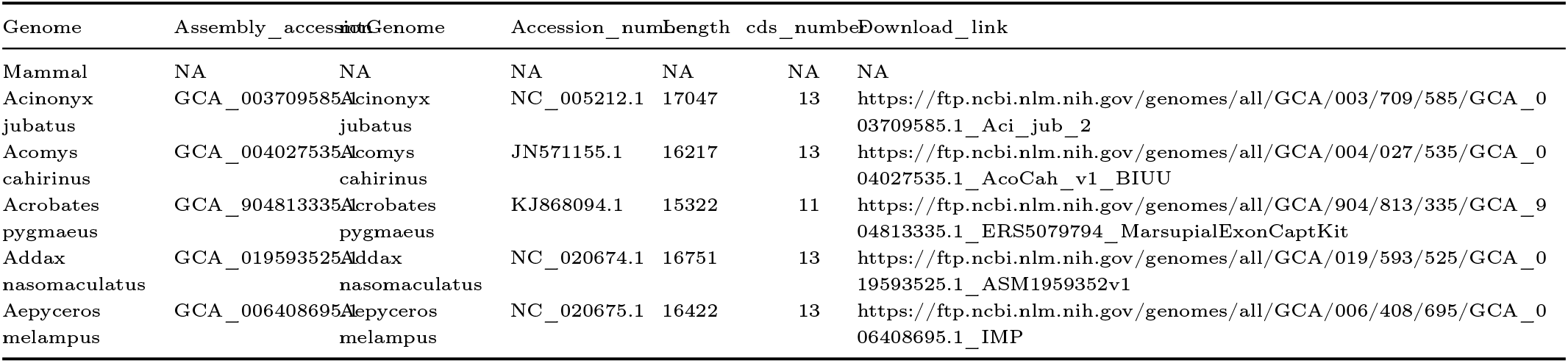
All identifiers of the genomes and mitogenomes and their respective species used in this study. For full table see Supplementary_tables.xlsx

## Notes

### Competing Interest Statement

The authors have declared no competing interest.

### Summary of Updates

The title has been changed to highlight both aspects (introgression and selection). A simulation section was added to the manuscript to investigate the heterogenous mutation landscape of nuclear and mitochondrial genome.

